# Risk aversion in macaques in a freely moving patch-leaving foraging task

**DOI:** 10.1101/452151

**Authors:** B. R Eisenreich, Benjamin Y. Hayden

## Abstract

Animals, including humans, are risk-averse in most contexts. A major exception is the rhesus macaque (*Macaca mulatta*), which is robustly risk-seeking. Macaques‘ unique preferences may reflect their unique evolutionary history. Alternatively, they may derive from elements of task design associated with the demands of physiological recording, the source of nearly all macaque risk preference data. To disambiguate these possibilities we assessed macaques’ risk attitudes in a somewhat more naturalistic environment: subjects foraged at four feeding stations in a large enclosure. Stations (i.e. patches) provided either stochastically or non-stochastically depleting rewards. Subjects’ patch residence times were longer at safe than at risky stations, indicating a preference for safe options. This preference was not attributable to a win-stay-lose-shift heuristic. These findings highlight the lability of risk attitudes in macaques and support the hypothesis that observed differences between macaques and other species are ephemeral, not evolved.

## INTRODUCTION

Many animals, including humans, prefer sure things to gambles (Kacelnik & Bateson, 1996). The tendency to minimize risk (i.e. unknowable and unpredictable variation) has been a topic of interest from behavioral ecology (Heilbronner, 2017; Stephens & Krebs, 1986) to economics (Kanheman & Tversky, 1979; O’Donoghue & Somerville, 2018) and neuroscience (Genest et al., 2016; Knutson & Bossaerts, 2007; Mccoy & Platt, 2005; Preuschoff et al., 2008; Strait et al., 2014; Calhoun and Hayden, 2015). Furthermore, cognitive processes related to decision making in risky contexts underlies many maladaptive behaviors such as addiction and problem gambling (Peters et al., 2016; Wilson & Vassileva, 2018). Consequently accurately understanding risk attitudes provides important insight into the evolutionary origin, and thus the psychological and neural mechanisms, of addiction and maladaptive choice (Rosati and Santos Annual Reviews).

Most animals are risk averse (Kacelnik & Bateson, 1996). For example, bumble bees, rats, starlings, and humans all demonstrate reliable risk aversion in standard experimental tasks (Kagel et al., 1986; Marsh & Kacelnik, 2002; Real, 1990; Weber et al., 2004). Other risk-averse animals include fish, pigeons, juncos, warblers, jays, and shrews (Kacelnik & Bateson, 1996). In marked contrast, rhesus macaques are robustly risk-seeking (Genest et al., 2016; Hayden & Platt, 2007; Mccoy & Platt, 2005; O’Neill & Schultz, 2010; So & Stuphorn, 2010; Stauffer et al., 2015; Xu & Kralik, 2014). Macaques’ exceptional risk preferences represent an abnormality under many theories of decision-making. For example, they are not consistent with the presumptions of prospect theory, probably the most widely used behavioral economic approaches to studying risk (Kanheman & Tversky, 1979). Meanwhile, scalar utility theory, which is based on granular psychological principles, predicts risk aversion due to inherent noise in memory processes for rewards (Kacelnik & Abreu, 1998). Even risk sensitive foraging theory (Caraco, 1981), which predicts risk-seeking as an adaptive strategy to energetically poor environments and deprived physiological states, does not fully explain macaque risk attitudes as experimental deprivation increases greater risk aversion in contradiction of the prediction of increased risk-seeking (Yamada et al. 2013).

Explanations for why macaques differ from other animals generally come in two types. One type of explanation assumes that macaques’ risk attitudes are an evolved reflection of their foraging history. Evolutionary factors that could promote risk-seeking in macaques include their specific feeding ecology and social structure (Heilbronner et al., 2008; Stevens et al., 2005; Santos & Rosati, 2015). For example, rhesus macaques forage in social groups on a wide range of herbaceous resources including human crops, as well as insects (Richard et al., 1989). Their broad diet allows for greater tolerance of variability within a resource due to many readily available alternatives. This tolerance for variability may reflect in their tolerance for variable options in economic tasks.

Another possibility is that macaques‘ risk-seeking is a measurement artifact. The manner in which macaques’ risk attitudes are measured is generally different from methods used for other species (Hayden & Platt, 2009; Heilbronner, 2017; Heilbronner & Hayden, 2013). Macaques are a popular model organism for neurophysiological studies of decision-making. Their risk attitudes tend to be assessed in contexts tailored to the needs of electrophysiology, not cross-species comparison. Thus they are tested with rapid trials (often as fast as three seconds per trial), small stakes, abstract stimuli, immediate rewards, overtraining, oculomotor responses, and hundreds or thousands of trials in a few hours. It may be that one of these factors, or some combination thereof, motivate risky choice. Indeed, even humans can become risk-seeking when gambling for small rewards in conditions designed to be similar to those used in non-human primate experiments (Hayden & Platt, 2009). And even modest changes, like slowing the pace of decisions by inserting inter-trial intervals, markedly reduces (but does not eliminate) risk-seeking in macaques (Hayden & Platt, 2007). Likewise, using token rewards alters other patterns of risk preferences, although it does not eliminate risk-seeking (Farashahi et al., 2018).

We hypothesized that with greater effort to make decisions more natural, we could make macaques risk-averse, and thus match behavior found in other species. We designed a naturalistic foraging task based on the patch-leaving problem from foraging theory (Stephens & Krebs, 1986; Charnov 1976; Nonacs, 2001). We tested subjects (n=3) using a single subject design within a large enclosure that allowed for free movement between different feeding stations. Our task design incorporates risk within the stochasticity of patch harvests rates (consequently, risk is orthogonal to reward amount). We allowed for self-governed allocation of behavior among patches. Thus, we are able to examine the influence of risk across the use of patch types in addition to within particular patches. We found that macaques are risk-averse under these foraging conditions. Two of the same subjects exhibited risk-seeking in a standard risk task designed for physiological recording, indicating that their risk preferences are task-specific, not individual-specific. Taken together, our results demonstrate the impact of the environmental structure on the malleable nature of risk attitudes in rhesus macaques and highlight the importance of using naturalistic tasks for studying cognitive processes.

## RESULTS

### The freely moving patch-leaving task

We examined responses of three macaque subjects in a novel *freely moving patch-leaving task* (see **Methods**). We defined the optimal harvest rate as the maximum of the gain function. The gain function is defined by the cumulative reward harvested divided by the total time spent foraging (Charnov, 1976) (figure 1). The optimal harvest is the rate that maximizes the gain function and serves as an index of when a forager should leave a patch. Across patch types the average leave time of subjects produced harvest rates close to the optimal. Had they left patches with no behavioral variability they would have obtained 90% of the reward an optimal forager would have (subject C= 93%, subject K= 89.19%, subject Y= 87.52%). The variability in patch leaving was around 4 turns (C: sd= 3.4 turns, K: sd= 4.5 turns, Y: sd= 4.8 turns,). All three exhibited a modest tendency to overharvest, meaning that they overstayed compared to an optimal forager, within a patch by 2-3 turns (average harvest lengths C: 8.636 turns, K: 9.3989 turns, and Y: 10.1599 turns, optimal harvest length 7 turns).

**Figure 1.**
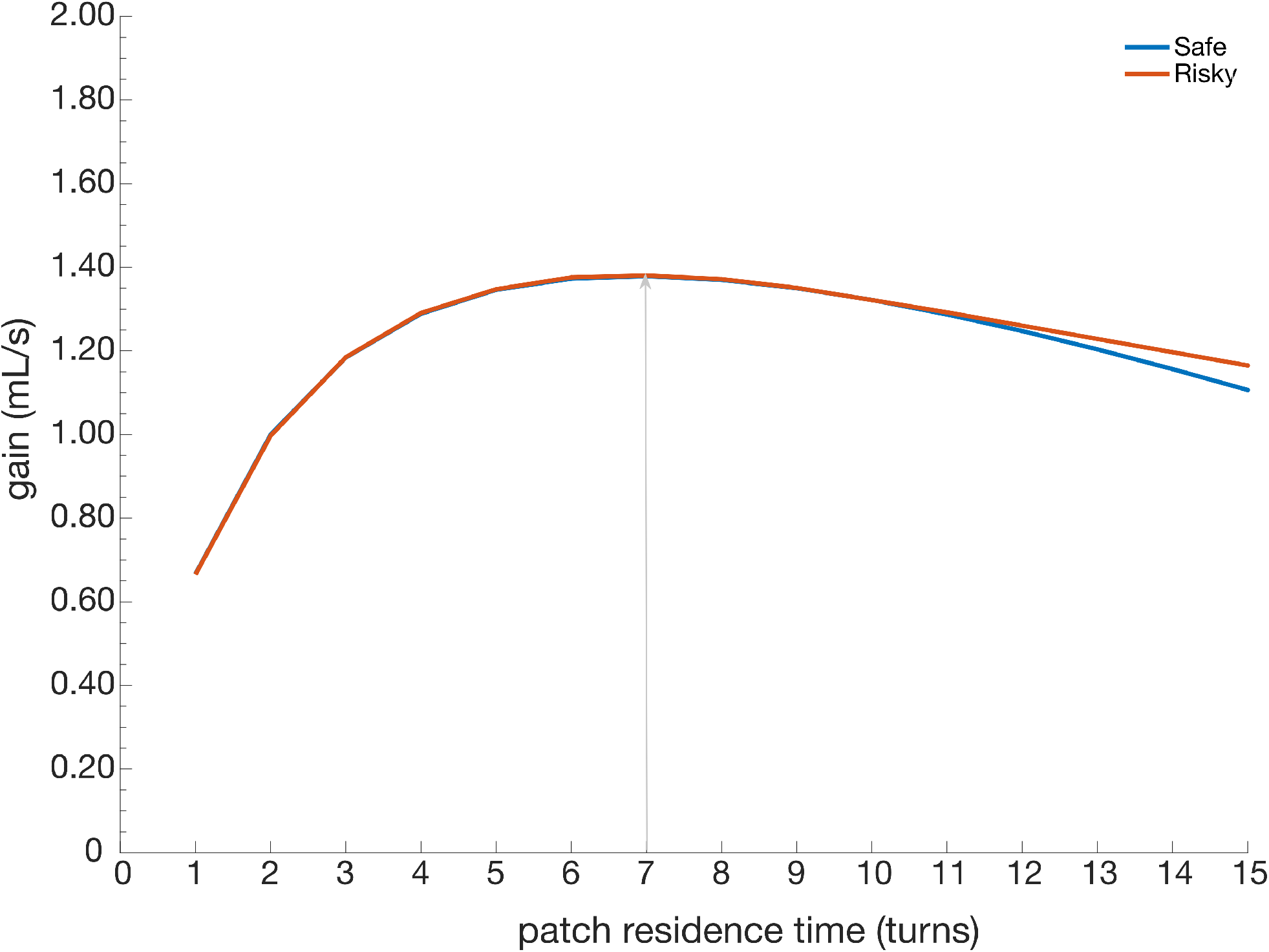
Gain function (rate as a function of residence time) for safe patches (blue line) and risky patches (red line). The black arrow denotes the abscissa point of the maximum intake rate, and thus the rate-maximizing strategy for both patch types. Due to the programed variation in reward amounts, the gain function for risky patches diverges slightly from the safe patch at long residence times. This divergence arises due to the limitation of reward amounts being bounded at 0 seconds of solenoid open time.

### Macaques are indifferent between choice of patch types

Within our foraging task, subjects freely forage within any of the four patches. Thus we can examine how risk influences both the choice of which patch to forage from, in addition to the decision of when to leave a patch. How risk impacts the choice of a patch has been largely unstudied. In our task all four patches have identical mathematical expected value (EV), such that a rate maximizing forager should show no preference between patch types. However a patch‘s subjective value may be influenced by the presence of risk, such that a forager would prefer one patch type to the other. Given macaques’ robust risk-seekingness, we would expect subjects to forage from risky patches more often than safe patches. Contrary to this prediction, we found no evidence to support a preference for either patch type indicative of subjects basing the choice of patch on average reward rates (C: t(249)=-0.2525, p=0.8009, K: t(177)=-0.899, p=0.3699, Y: t(185)=-1.0267, p=0.3059).

### Macaques spend more time in safe patches than risky ones

Foragers calibrate patch residence time to the harvest rate within a patch (Charnov, 1976; Stephens & Krebs, 1986). Their residence time then provides a measure of the subjective value they assign to marginal rewards. This arises out of the putative decision variable that contrasts the marginal gain of staying within the patch against the average rate of reward gained from foraging within the environment. For our task, risk likely influences the subjective evaluation of marginal rewards within risky patches, as on each harvest the subject has no knowledge of whether the reward schedule will step up or down and thus has a noisy estimate of the local patch harvest rate.

We next examined residence time in safe and risky patches. All three subjects remained in the safe patches longer than in the risky ones (figure 2) (C: 0.9479 turns, t(248)=2.198, p=0.0144, d=0.278; K: 1.35 turns, t(176)= 2.0289, p=0.022, d= 0.304; Y: 1.17 turns, t(184)= 1.6842, p=0.0469, d=0.247). Furthermore, subjects stayed longer in safe patches despite it being suboptimal to do so (specifically, they obtained on average 88.71% of the total harvest rate opposed to 93.15%; C: 91.92% vs. 95.56%, K: 88.38% vs. 93.56%, Y: 85.82 % vs. 90.32%). The tendency of all three subjects to leave risky patches earlier resulted in more optimal harvesting and contrasts with reported biases for overstaying in patch tasks (Nonacs, 2001; Blanchard & Hayden, 2015). The apparent contrast between optimality and patch staying strongly supports the notion of subjective valuations in favor of avoiding risk as a key component in the decision to leave a patch.

**Figure 2.**
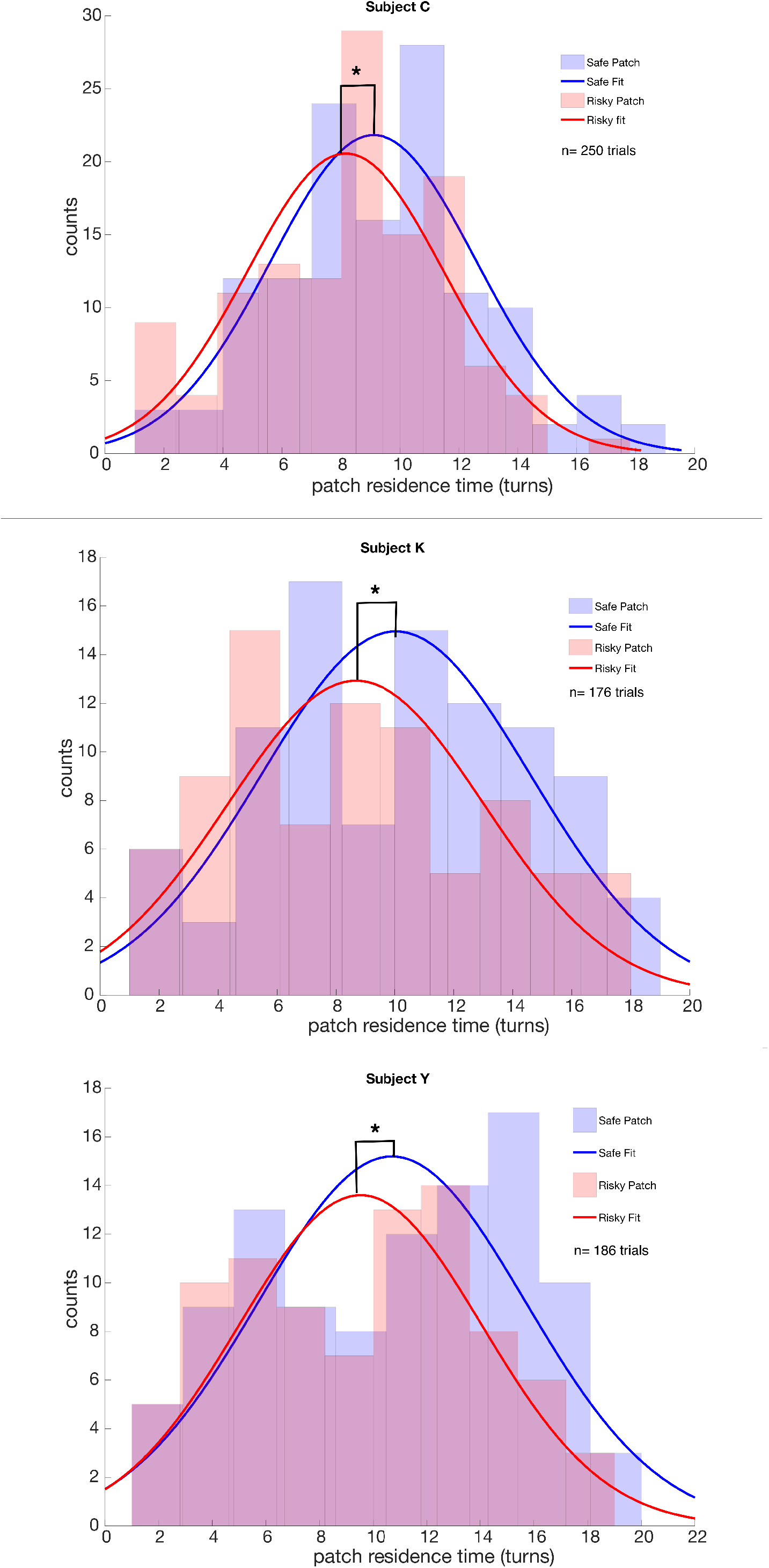
Histogram of residence times for all subjects in safe (blue) and risky (red) patches within the standard environment. Solid lines indicate Gaussian fits to the observed leaving times. Residence times are significantly longer for safe than risky patches, indicating risk aversion.

### No evidence for win-stay/loose-shift heuristic in guiding patch-leaving

It is possible that macaques’ longer residence times in safe patches is due to a data censoring effect: perhaps they leave when any individual outcome is lower than some threshold. That is, they may obey a win-stay loose-shift heuristic (Hayden et al., 2008; Hayden et al., 2009; Barraclough et al., 2004; Seo & Lee, 2007). To determine if subjects used this heuristic, we examined the likelihood of leaving a risky patch given the recent history of wins and losses. Even when combining across all sessions, none of the three subjects exhibited a significant preference of increased patch-leaving immediately after losses (C: t(122)=1.1740, p=0.2427, K: t(82)= 0.5465, p=0.5862, Y: t(85)= 0.6448, p=.05208). Nor did we observe any effect of harvest outcomes two steps back (C: F(3,119)=0.83, p=0.8009, K: F(3,79)=0.13, p=0.9413; Y: F(3,82)=1.44, p=0.237).

### Risk preferences shift with the coefficient of variation

Given macaques’ reported preference for risk we may expect that there is a unique species level difference in how risk is evaluated in the decision making process. Risk preferences may be governed by the experienced variance in reward or by the coefficient of variation (Ludvig et al., 2014; Weber et al., 2004, Shafir, 2000). In our task the experienced variance is the deviation in reward delivery times (in this case +/- 1s solenoid open time). The coefficient of variation is then the deviation in reward delivery divided by the average reward. We investigated whether subjects risk preferences were governed by changes t0 the coefficient of variation by increasing the magnitude of the reward schedule for both feeder types (risky and safe) while keeping variance the same. Increasing the reward magnitude while maintaining variance reduces the coefficient of variation (CV(standard)= 0.5 vs. CV(rich)=0.25) and has been demonstrated to produce less risk aversion in humans, pigeons, rats, and bees (Ludvig et al., 2014; Weber et al., 2004, Shafir, 2000). By contrast, if subject’s risk preferences are influenced by the variance there should be no change in behavior. In all three subjects we saw shifts away from risk-aversion to risk-neutrality/seeking as the coefficient of variation decreased (figure 3.) (C: t(117)= 3.3303, p=0.0005, d=0.605, K: t(99)=1.2077, p=0.115, d=0.226, Y: t(94)=1.7483, p=0.0418, d=0.351).

**Figure 3.**
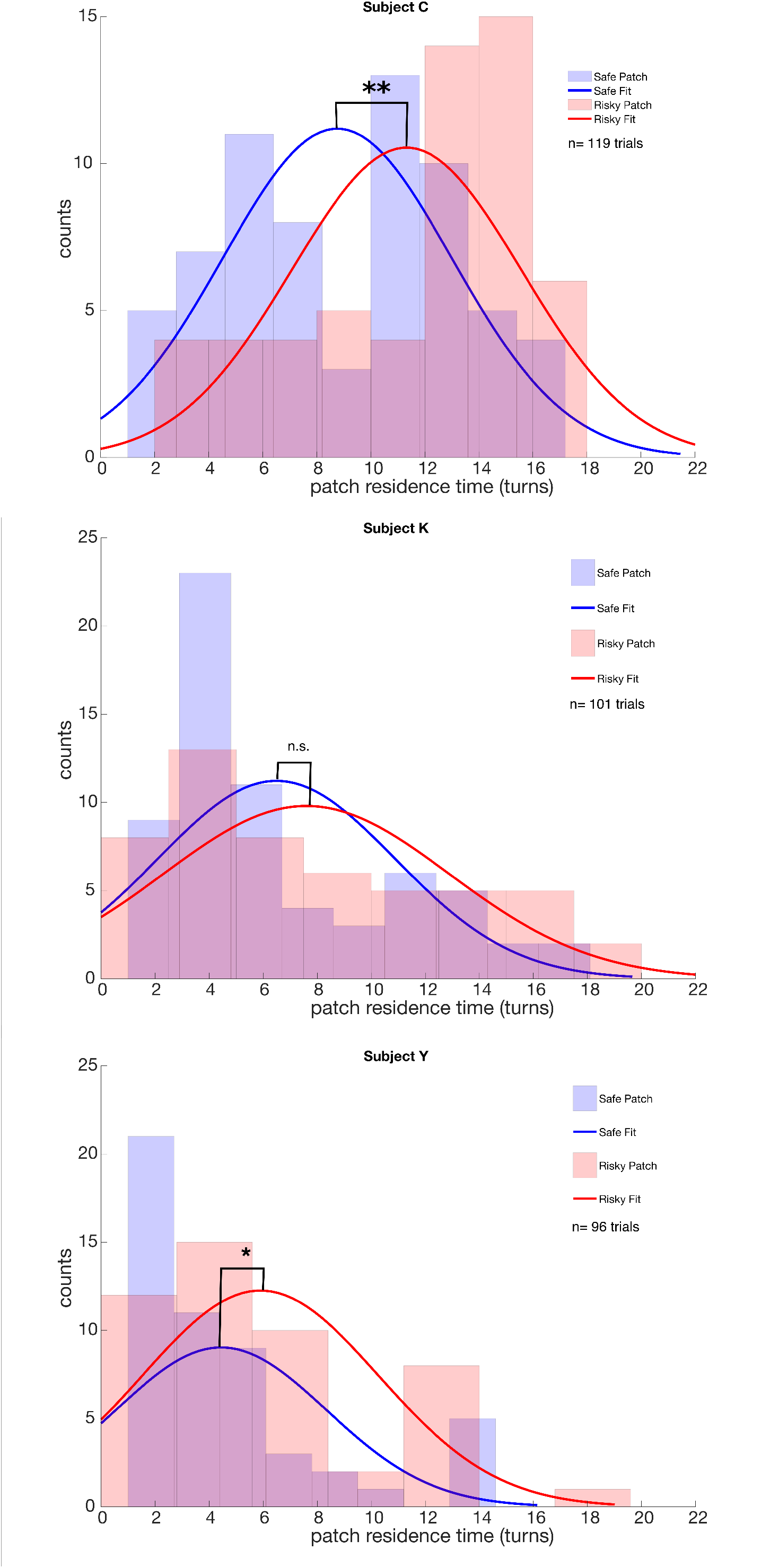
Histogram of residence time for all subjects in rich environment version of task. Plots follow the same conventions as figure 2. Subjects resided longer in risky patches than safe patches when the entire reward schedule for all feeder types was increased while maintaining the same variance as used in the standard environment.

### Two of the same macaques are risk-prone in computerized task

We next analyzed risky choice behavior in two subjects (C and K) in a standard (not foraging-based, not freely moving) juice gambling task (Strait et al., 2014). Both subjects exhibited strong risk-seeking behavior. On trials with matched expected values subject C choose the risky option 67% (t(1232)= 12.86, p<0.0001) of the time, while subject K choose the risky option 66% of the time (t(1437)= 12.55, p<0.0001).

This preference can be quantified using the shape of the utility curve. Both subjects showed convex utility curves (figure 4, C: alpha= 2.284, 95%CI = 2.584-1.983; K: alpha= 3.632, 95%CI=3.822-3.441). However within the more naturalistic risky patch task the same subjects exhibited concave utility curves indicative of strong risk aversion (figure 3, C: alpha= 0.550, 95% CI= 0.5922-0.508; K: alpha=0.743, 95%CI= 0.889-0.586).

**Figure 4.**
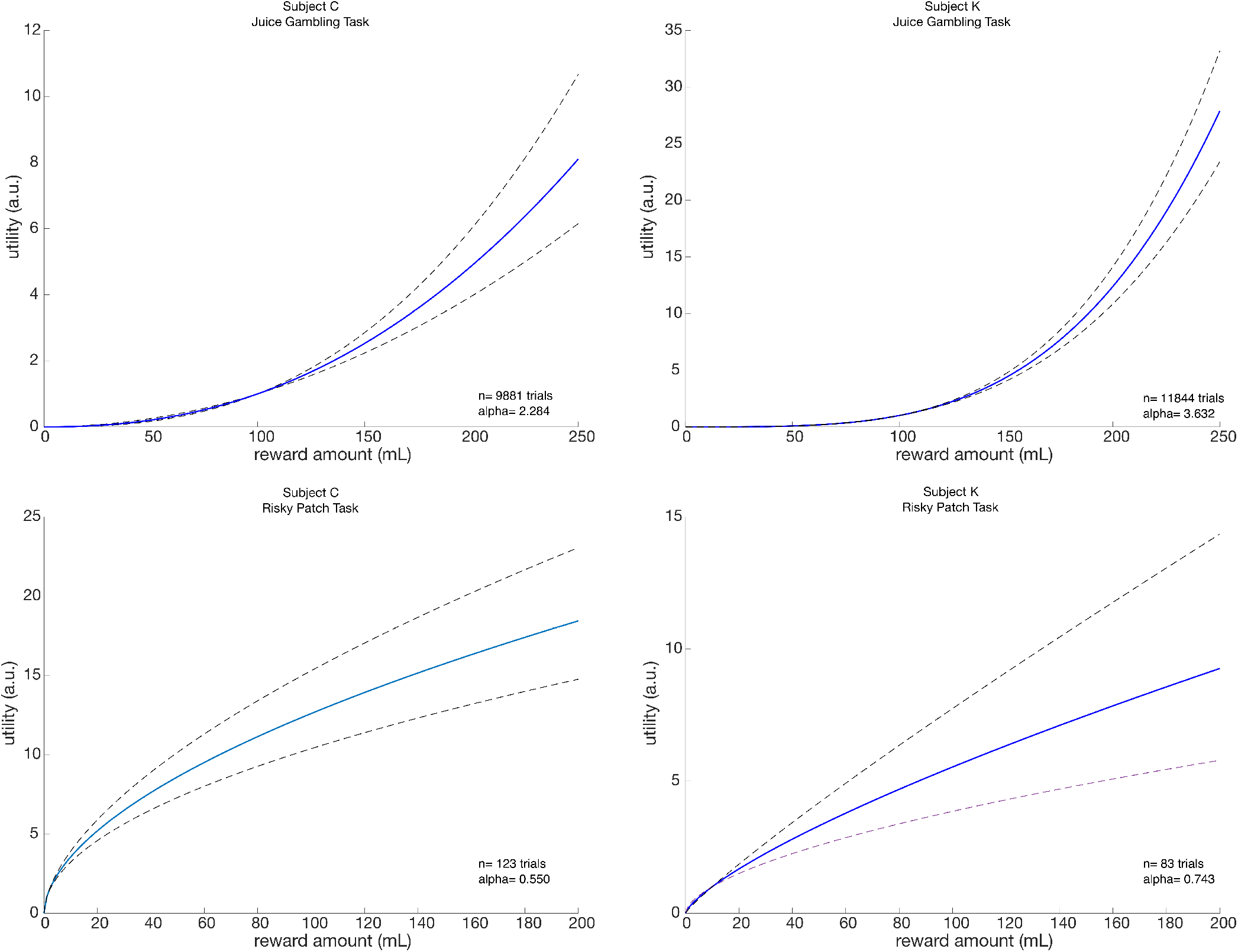
Plotted utility functions for two subjects who participated in both the feely moving patch task (lower panels) and a standard chaired economic task (upper panels). Dotted lines represent 95% CI. Two of the same macaques are risk-seeking in the standard task (convex utility curves), and risk-averse the freely moving patch task (concave utility curves).

## DISCUSSION

Risk is ubiquitous and unavoidable in the natural environment. Foragers must have strategies for dealing with it. Macaques are found to be risk-seeking in many labs and many contexts (Mccoy & Platt, 2005) Blanchard, Wilke, and Hayden, 2014; Heilbronner and Hayden, PBR, 2015; (Genest et al., 2016; O’Neill & Schultz, 2010; Xu & Kralik, 2014). Risk-seeking in macaques is not due to a preference for variability nor can it be explained by the shape of the utility curve (Heilbronner et al., 2009). It exists alongside a robust ambiguity aversion (Hayden, Heilbronner, & Platt, 2010), and is partially but not fully explainable as a motivation for information (Blanchard, Hayden, & Bromberg-Martin, 2015); Kidd and Hayden, 2016).

Macaques’ risk-seeking is puzzling because most animals are risk-averse. Indeed, a major meta-analysis shows robust risk aversion across dozens of taxa (Kacelnik & Bateson, 1996). Likewise, humans have been shown to be risk averse (at least for gains) in a large number of cases (Holt and Laury, 2002; Rabin and Thaler, JEP, 2001). Within this, context, the reliable risk-seeking patterns of rhesus macaques presents an important challenge. One approach to reconciling this inconsistency – and more generally, any dissimilarity between species – is to consider their unique evolutionary history, especially their foraging history (Heilbronner et al., 2008; Santos & Rosati, 2015; Stephens, 2008; Todd & Gigerenzer, 2007). An alternative approach is to abandon the idea that simple economic tasks measure a robust and invariable trait (Farashahi et al., 2018).

This alternative viewpoint starts with the observation that foragers, especially primates, are highly cognitively flexible. They can adapt their strategies rapidly and gracefully to changing circumstances to optimize their intake, perhaps with unobservable constraints. There is some evidence that macaque risk-seeking is labile: (1) changing the delays between trials can increase or reduce risk-seeking, although it does not produce risk aversion (Hayden & Platt, 2007); (2) across ostensibly similar tasks, very different patterns of risk attitudes are observed, even as those patterns are consistent between subjects (Farashahi et al., 2018); and (3) when humans are tested for risk attitude in envionments designed to mimic those that produce risk-seeking in macaques, they show trends towards risk-seeking as well (Hayden & Platt, 2009). Finally, other studies indicate that apparent impulsivity in macaques may be partially attributable to experimental confounds (Hayden, PBR, 2016; Pearson, Hayden, and Platt, Frontiers, 2010).

These points point towards the possibility that there is something special about the tasks used to measure risk attitudes in macaques that favors risk-seeking behavior (Heilbronner & Hayden, 2013). To test this idea, we measured risk attitudes in macaques in a more naturalistic context. We embedded the decisions in the context of a patch-leaving task and implemented them in a large space around which macaques could move freely and unconstrained. In this context, we found reliable risk-aversion. Contrary to species level explanations, our results implicate the inherent structure of behavioral tasks as the primary driver of macaque risk preference. This does not mean that species differences have no role in risk attitudes. Instead, they suggest that risk attitudes are so labile that one must carefully consider all parameters of task design when interpreting economic preferences (Stephens and Anderson, 2001). More fundamentally, these results suggest that animals may not have a stable risk attitude, but rather have a consistent but flexible cognitive repertoire that they use when encountering risk. Future studies will be needed to tabulate an inventory of this repertoire.

## METHODS

### Subjects and Apparatus

Three male rhesus macaques served as subjects for the experiment. Two of the subjects (C and K) had previously served as subjects on standard neuroeconomic tasks, including a set shifting task (Sleezer and Hayden, 2016), a diet selection task (Blanchard and Hayden, 2014; Blanchard and Hayden, 2015), and intertemporal choice tasks (Blanchard, Pearson, & Hayden, 2013). while the third subject (Y) was naïve to all experimental procedures. All three monkeys were fed ad libitum and pair housed within a light and temperature controlled colony room. All research and animal care was conducted in accordance with university IACUC approval and in accord with NIH standards for the care and use of non-human primates.

Subjects were behaviorally tested in a large cage (8‘ L × 9’ W × 8’H) made framed panels consisting of 2 inch wire mesh. This allowed for free movement of the subjects within the cage in three dimensions. Five 55 gallon drum barrels weighted with sand were placed within the cage to serve as perches for the subjects to sit upon. Four juice feeders were placed at each of the four corners of the cage in a rotationally symmetric alignment. The juice feeders consisted of a 16×16 led screen, a lever, buzzer, a solenoid (Parker Instruments), and an arduino uno. Data was collected in MatLab (Mathworks) via Bluetooth communication with each of the juice feeders.

### Behavioral Tasks

#### Patch leaving risk task

The patch leaving risk task is designed to mimic the natural depletion of prey items from a patch the longer a subject forages from it (cf. Hayden, Pearson, and Platt, 2011; Blanchard and Hayden, 2015). Each feeder was programed to deliver a base reward schedule consisting of an initial 2 sec of juice that decreased by 0.125 sec with each subsequent delivery (turn) (standard environment). In the rich environment, the feeders provided 4 sec of juice that decreased by 0.25 sec each turn. Risk, here defined as variation in reward times, was introduced by programming two of the juice feeders to randomly increase or decrease the juice delivery time by 1 sec in addition to the base reward schedule at a probability of 0.5. Both feeder types delivered rewards following their respective schedules until reaching the base value of 0, at which point the patch is depleted and no more rewards were delivered. In practice this depletion process results in identical gain functions over the majority of patch residence times. However because the schedule had a bound at 0 seconds, the tail end of the gain function for risky patches does diverge from safe patches (**Figure 1**).

Two of the four feeders diagonally across from each other were designated as variable feeders, while the other two served as safe feeders (no variation in reward delivery). Feeders were visually identical, although they could be readily discriminated by their position relative to landmarks outside the cage. The feeder designations remained fixed for each subject across experimental days. Each feeder displayed the total amount of prey available within the patch via a blue bar (8×16 LEDs). With each lever press, juice would be delivered and a portion of the blue bar would disappear, explicitly indicating its depletion status. Leaving a feeder to activate any of the other three feeders would cause the previously activated feeder to immediately fully replenish.

#### Juice Gambling task

The juice gambling task, which we used as a comparison, was used previously collected for electrophysiology experiments (Strait et al. 2014; Strait et al., 2015a; Strait et al., 2015b). In brief the task consisted of paired choices presented rapidly (~3 sec duty cycle) while subjects sat in a chair. Choices were made rapidly with saccades to spots on a computer screen. Stimuli were colored bars that indicated probability and stakes.

### Data Analysis

#### Patch leaving risk task

Behavioral data were analyzed in Matlab. For the patch leaving risk task we defined the duration of patch residence as the number of presses at a given juice feeder. Data were combined across experimental days for each subject. In order to analyze the overall optimality of subjects within the foraging task, we used the marginal value theorem (Charnov, 1976) to determine the optimal leaving time.

#### Risk Parameter Estimation

To analyze differences in risk preferences between the juice gambling task and our patch leaving risk task we fit each subject’s choice preferences for offer 1 or for the decision to stay in the current patch to the two equations below (Eq. 1 and Eq. 2) using maximum likelihood estimation. In both equations the parameter a functions as an index of risk preference such that a< 1 implies greater risk avoidance, a>1 risk-seeking, and a=1 risk neutrality.

*Eq. 1 Juice Gambling Task*

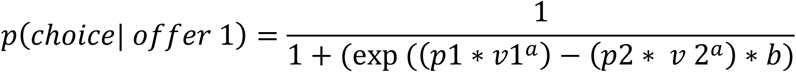

Where:

p1= probability of offer 1

v1= value of offer 1 (s)

p2= probability of offer 2

v2= value of offer 2(s)

a= risk preference index

b= measure of stochasticity

*Eq. 2 Patch Leaving Risk Task*

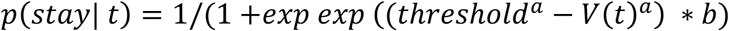

Where:

t= time measured in discrete lever presses

threshold= point of indifference between staying and leaving a patch

V(t)= current reward amount available given the time spent in the patch

a= risk preference index

b= measure of stochasticity

## References

Agetsuma, N (1995a) Foraging strategies of Yakushima macaques *(Macaca fuscata yakui)*. Int J Primatol 16:595–609.

Barraclough, D. J., Conroy, M.L., & Lee, D. (2004). Prefrontal cortex and decision making in a mixed-strategy game. Nature Neuroscience, 7, 404–410. doi: 10.1038/nm1209.

Blanchard, T. C, & Hayden, B. Y. (2014). Neurons in dorsal anterior cingulate cortex signal postdecisional variables in a foraging task. Journal of Neuroscience, 34(2), 646–655.

Blanchard, T. C, & Hayden, B. Y. (2015). Monkeys Are More Patient in a Foraging Task than in a Standard Intertemporal Choice Task. PLoS ONE, 1-11. http://doi.org/10.1371/journal.pone.0117057.

Blanchard, T. C, & Hayden, B. Y. (2015). Ramping ensemble activity in dorsal anterior cingulate neurons during persistent commitment to a decision. Journal of Neurophysiology, 114(4), 2439–2449.

Blanchard, T. C, Hayden, B. Y., & Bromberg-Martin, E. S. (2015). Orbitofrontal cortex uses distinct codes for different choice attributes in decisions motivated by curiosity. Neuron, 85(3), 602–614. http://doi.org/10.1016/j.neuron.2014.12.050.

Blanchard, T. C, Pearson, J. M., & Hayden, B. Y. (2013). Postreward delays and systematic biases in measures of animal temporal discounting. Proceedings of the National Academy of Sciences, 201310446.

Caraco, T. (1981). Energy budgets, risk and foraging preferences in dark-eyed juncos (Junco hyemalis). Behavioral Ecology and Sociobiology, 8(3), 213–217. http://doi.org/10.1007/BF00299833.

Charnov, E. L. (1976). Optimal Foraging, the Marginal Value Theorem. Theoretical Population Biology, 9(2), 129–136.

De Petrillo, F., Ventricelli, M., Ponsi, G., & Addessi, E. (2015). Do tufted capuchin monkeys play the odds? Flexible risk preferences in Sapajus spp. Animal Cognition, 18(1), 119–130. http://doi.org/10.1007/s10071-014-0783-7.

Farashahi, S., Azab, H., Hayden, B., & Soltani, A. (2018). On the Flexibility of Basic Risk Attitudes in Monkeys. The Journal of Neuroscience, 38(18), 4383–4398. http://doi.org/10.1523/JNEUROSCI.2260-17.2018.

Genest, W., Stauffer, W. R., & Schultz, W. (2016). Utility functions predict variance and skewness risk preferences in monkeys. Proceedings of the National Academy of Sciences, 113(30). http://doi.org/10.1073/pnas.1602217113.

Hayden, B. Y. (2018). Economic choice: the foraging perspective. Current Opinion in Behavioral Sciences, 24, 1-6. http://doi.org/10.1016/J.COBEHA.2017.12.002.

Hayden, B. Y., Heilbronner, S. R., & Platt, M. L. (2010). Ambiguity aversion in rhesus macaques. Frontiers in Neuroscience, 4(SEP), 1-7. http://doi.org/10.3389/fnins.2010.00166.

Hayden, B. Y., & Platt, M. L. (2007). Temporal Discounting Predicts Risk Sensitivity in Rhesus Macaques. Current Biology, 17(1), 49-53. http://doi.org/10.1016/j.cub.2006.10.055.

Hayden, B. Y., & Platt, M. L. (2009). Gambling for Gatorade: risk-sensitive decision making for fluid rewards in humans. Animal Cognition, 12, 201–207. http://doi.org/10.1007/s10071-008-0186-8.

Hayden, B. Y., Pearson, J. M., & Platt, M. L. (2011). Neuronal basis of sequential foraging decisions in a patchy environment. Nature Neuroscience, 14(7), 933.

Heilbronner, S. R. (2017). Modeling risky decision-making in nonhuman animals舁: shared core features. Current Opinion in Behavioral Sciences, 16, 23–29. http://doi.org/10.1016/j.cobeha.2017.03.001.

Heilbronner, S. R., & Hayden, B. Y. (2013). Contextual factors explain risk-seeking preferences in rhesus monkeys. Frontiers in Neuroscience, 7(7 FEB), 1–7. http://doi.org/10.3389/fnins.2013.00007.

Heilbronner, S. R., Rosati, A. G., Stevens, J. R., Hare, B., & Hauser, M. D. (2008). A fruit in the hand or two in the bush舁? Divergent risk preferences in chimpanzees and bonobos. Biology Letters, 4(March), 246–249. http://doi.org/10.1098/rsbl.2008.0081.

Kacelnik, A., & Abreu, F. B. e. (1998). Risky Choice and Weber’s Law. Journal Theoretical Biology, 194(jt980763), 289–298.

Kacelnik, A., & Bateson, M. (1996). Risky Theories — The Effects of Variance on Foraging Decisions 1. American Zoologist, 434, 402–434.

Kagel, J. H., MacDonald, D. N., Battalio, R. C., White, S., & Green, L. (1986). Risk aversion in rats (Rattus norvegicus) under varying levels of resource availability. Journal of Comparative Psychology, 100(2), 95–100. http://doi.org/10.1037/0735-7036.100.2.95.

Kanheman, D., & Tversky, A.. (1979). Prospect theory: An analysis of decision under risk. Econometrica, 47, 263–291.

Knutson, B., & Bossaerts, P. (2007). Neural Antecedents of Financial Decisions. Journal of Neuroscience, 27(31), 8174–8177. http://doi.org/10.1523/JNEUROSCI.1564-07.2007.

Ludvig, E. A., Madan, C. R., Pisklak, J. M., & Spetch, M. L. (2014). Reward context determines risky choice in pigeons and humans. Biology Letters, 10(8). http://doi.org/10.1098/rsbl.2014.0451.

Marsh, B., & Kacelnik, A. (2002). Framing effects and risky decisions in starlings. Proceedings of the National Academy of Sciences, 99(5), 3352–3355.

Mccoy, A. N., & Platt, M. L. (2005). Risk-sensitive neurons in macaque posterior cingulate cortex. Nature Neuroscience, 8(9), 1220–1227. http://doi.org/10.1038/nn1523.

Nonacs, P. (2001). State dependent behavior and the marginal value theorem. Behavioral Ecology, 12(1), 71-83.

O’Donoghue, T., & Somerville, J. (2018). Modeling Risk Aversion in Economics. Journal of Economic Perspectives, 32(2), 91–114. http://doi.org/10.1257/jep.32.2.91.

O’Neill, M., & Schultz, W. (2010). Coding of reward risk by orbitofrontal neurons is mostly distinct from coding of reward value. Neuron, 68(4), 789–800. http://doi.org/10.1016/j.neuron.2010.09.031.

Peters, S. K., Dunlop, K., & Downar, J. (2016). Cortico-Striatal-Thalamic Loop Circuits of the Salience Network舁: A Central Pathway in Psychiatric Disease and Treatment. Frontiers in Systems Neuroscience, 10(December), 1–23. http://doi.org/10.3389/fnsys.2016.00104.

Preuschoff, K., Quartz, S. R., & Bossaerts, P. (2008). Human Insula Activation Reflects Risk Prediction Errors As Well As Risk. Journal of Neuroscience, 28(11), 2745–2752. http://doi.org/10.1523/JNEUROSCI.4286-07.2008.

Proctor, D., Williamson, R. A., Latzman, R. D., de Waal, F. B. M., & Brosnan, S. F. (2014). Gambling primates: Reactions to a modified Iowa Gambling Task in humans, chimpanzees and capuchin monkeys. Animal Cognition, 17(4), 983–995. http://doi.org/10.1007/s10071-014-0730-7.

Real, L. A. (1990). Search Theory and Mate Choice. I. Models of Single-Sex Discrimination Author(s): Leslie Real Source舁: The American Naturalist, Vol. 136, No. 3 (Sep., 1990), pp. 376-405 Published by舁: The University of Chicago Press for The American Society o. The American Naturalist, 136(3), 376–405.

Santos, L. R., & Rosati, A. G. (2015). The Evolutionary Roots of Human Decision Making. Annual Reviews Psychology, 66, 321–347. http://doi.org/10.1146/annurev-psych-010814-015310.

The Sayers, K., & Menzel, C. R. (2017). Risk sensitivity, phylogenetic reconstruction, and four chimpanzees. Behavioral Ecology and Sociobiology, 71(1). http://doi.org/10.1007/s00265-016-2234-8.

Seo, H., & Lee, D. (2007). Temporal filtering of reward signals in the dorsal anterior cingulate cortex during a mixed strategy game. Journal of Neuroscience, 27(31), 8366–8377.

Shafir, S. (2000). Risk-sensitive foraging: The effect of relative variability. Oikos, 88, 663–669.

Sleezer, B.J., & Hayden, B.Y. (2016). Differential contributions of ventral and dorsal striatum to early and late phases of cognitive set reconfiguration. Journal of Cognitive Neuroscience, 28(12), 1849–1864.

So, N.-Y., & Stuphorn, V. (2010). Supplementary Eye Field Encodes Option and Action Value for Saccades With Variable Reward. Journal of Neurophysiology, 104(5), 2634–2653. http://doi.org/10.1152/jn.00430.2010.

Stauffer, X. W. R., Lak, X. A., Bossaerts, P., & Schultz, W. (2015). Economic Choices Reveal Probability Distortion in Macaque Monkeys. The Journal of Neuroscience, 35(7), 3146–3154. http://doi.org/10.1523/JNEUROSCI.3653-14.2015.

Stephens, D. W. (2008). Decision ecology舁: Foraging and the ecology of animal decision making. Cognitive Affective Behavioral Neuroscience, 8(4), 475–484. http://doi.org/10.3758/CABN.8.4.475.

Stephens, D. W., & Krebs, J. R. (1986). Foraging Theory. Princenton: Princenton University Press.

Stevens, J. R., Rosati, A. G., Ross, K. R., & Hauser, M. D. (2005). Will travel for food: Spatial discounting in two New World monkeys. Current Biology, 15(20), 1855–1860. http://doi.org/10.1016/j.cub.2005.09.016.

Strait, C. E., Blanchard, T. C., & Hayden, B. Y. (2014). Reward value comparison via mutual inhibition in ventromedial prefrontal cortex. Neuron, 82(6), 1357–1366. http://doi.org/10.1016/j.neuron.2014.04.032.

Strait, C. E., Sleezer, B. J., & Hayden, B. Y. (2015a). Signatures of value comparison in ventral striatum neurons. PLoS Biology, 13(6), e1002173.

Strait, C. E., Sleezer, B. J., Blanchard, T. C., Azab, H., Castagno, M. D., & Hayden, B. Y. (2015b). Neuronal selectivity for spatial positions of offers and choices in five reward regions. Journal of Neurophysiology, 115(3), 1098-1111.

Todd, P. M., & Gigerenzer, G. (2007). Environments That Make Us Smart. Current Directions in Psychological Science, 16(3), 167–171. http://doi.org/10.1111/j.1467-8721.2007.00497.x.

Weber, B. J., & Chapman, G. B. (2005). Playing for peanuts舁: Why is risk seeking more common for low-stakes gambles舁? Organizational Behavior and Human Decision Processes, 97, 31–46. http://doi.org/10.1016/j.obhdp.2005.03.001.

Weber, E. U., Shafir, S., & Blais, A. R. (2004). Predicting Risk Sensitivity in Humans and Lower Animals: Risk as Variance or Coefficient of Variation. Psychological Review, 111(2), 430–445. http://doi.org/10.1037/0033-295X.111.2.430.

Wilson, M. J., & Vassileva, J. (2018). Decision-making under risk, but not under ambiguity, predicts pathological gambling in discrete types of abstinent substance users. Frontiers in Psychiatry, 9(JUN), 1–10. http://doi.org/10.3389/fpsyt.2018.00239.

Xu, E. R., & Kralik, J. D. (2014). Risky business舁: rhesus monkeys exhibit persistent preferences for risky options. Frontiers in Psychology, 5(April), 1–12. http://doi.org/10.3389/fpsyg.2014.00258.

Yamada, H., Tymula, A., Louie, K., & Glimcher, P. W. (2013). Thirst-dependent risk preferences in monkeys identify a primitive form of wealth. Proceedings of the National Academy of Sciences, 110(39), 15788–15793. http://doi.org/10.1073/pnas.1308718110/-/DCSupplemental.www.pnas.org/cgi/doi/10.1073/pnas.1308718110.

